# Automated Segmentation of Fractured Distal Radii by 3D Geodesic Active Contouring of *in vivo* HR-pQCT Images

**DOI:** 10.1101/2020.10.14.339739

**Authors:** Nicholas Ohs, Caitlyn J. Collins, Duncan C. Tourolle, Penny R. Atkins, Bryant Schroeder, Michael Blauth, Patrik Christen, Ralph Müller

## Abstract

Radius fractures are among the most common fracture types; however, there is limited consensus on the standard of care. A better understanding of the fracture healing process could help to shape future treatment protocols and thus improve functional outcomes of patients. High-resolution peripheral quantitative computed tomography (HR-pQCT) allows monitoring and evaluation of the radius on the micro-structural level, which is crucial to our understanding of fracture healing. However, current radius fracture studies using HR-pQCT are limited by the lack of automated contouring routines, hence only including small number of patients due to the prohibitively time-consuming task of manually contouring HR-pQCT images.

In the present study, a new method to automatically contour images of distal radius fractures based on 3D morphological geodesic active contours (3D-GAC) is presented. Contours of 60 HR-pQCT images of fractured and conservatively treated radii spanning the healing process up to one year post-fracture are compared to the current gold standard, hand-drawn 2D contours, to assess the accuracy of the algorithm. Furthermore, robustness was established by applying the algorithm to HR-pQCT images of intact radii of 73 patients and comparing the resulting morphometric indices to the gold standard patient evaluation including a threshold- and dilation-based contouring approach. Reproducibility was evaluated using repeat scans of intact radii of 19 patients.

The new 3D-GAC approach offers contours within inter-operator variability for images of fractured distal radii (mean Dice score of 0.992 ± 0.004 versus median operator Dice score of 0.993 ± 0.006). The generated contours for images of intact radii yielded morphometric indices within the *in vivo* reproducibility limits compared to the current gold standard. Additionally, the 3D-GAC approach shows an improved robustness against failure (n = 4) when dealing with cortical interruptions, fracture fragments, etc. compared with the automatic, default manufacturer pipeline (n = 40). Using the 3D-GAC approach assures consistent results, while reducing the need for time-consuming hand-contouring.

## 1 Introduction

In our ageing population, bone fractures are increasingly common and have become a major socioeconomic burden [1]. Specifically, fractures of the distal radius are among the most common fracture types and indicative of reduced bone quality [2]. However, there is currently limited consensus on the optimal treatment protocol [3]. The conservative treatment of many radius fractures provides the opportunity to study the fracture healing process in humans. A better understanding of this process could help to shape future treatment protocols of distal radius fractures, ultimately resulting in better functional outcomes for patients.

With the introduction of high-resolution peripheral quantitative computed tomography (HR-pQCT), longitudinal changes in the microstructure of the radius can be monitored non-invasively [4–8]. Use of this technology has revealed critical changes in bone microstructure as a result of aging as well as pharmaceutical and surgical interventions. For example, an increase in cortical porosity, which is highly associated with a decrease in bone strength, was demonstrated in a longitudinal study of patients who had undergone kidney transplantation [8]. Preliminary evidence suggests that HR-pQCT might also be a viable technique to monitor bone fractures *in vivo* [9]. The microstructure in such studies is of fundamental importance, as techniques such as micro-finite-element analysis can be utilized to estimate the capability of a bone scanned with HR-pQCT to withstand mechanical load, a key parameter to assess successful fracture healing.

Image-based clinical research, such as that using HR-pQCT, requires accurate, reproducible and scalable tools for image processing. For this reason, automatic contouring approaches have been developed to reduce the need for hand-drawn contours [10–13] in HR-pQCT studies. The method developed by Buie et al. [10], specifically, is integrated into the software of the manufacturer of HR-pQCT devices (XtremeCT I and II, Scanco Medical AG, Switzerland), making it the de-facto standard. All of these automatic contouring approaches segment the target bone (i.e. radius) by first finding the outer contour of the bone with the assumption that the largest cortical interruptions imaged with HR-pQCT are small (on the sub-mm length scale) and that the cortex forms a well-defined, high-contrast edge in the HR-pQCT image. However, these assumptions break down when processing images of fractured radii where the cortex is often fragmented, and the presence of cartilaginous callus may blur the boundary between bone and soft tissue. Furthermore, radius fractures often occur in the ultra-distal region, contributing to the complexity of the image segmentation, as the cortex is both thinner and less mineralized. To our knowledge, approaches to generate automatic outer contours for HR-pQCT images of distal radius fractures have not been published yet.

Active contouring [14] is a promising technique that has already been shown to successfully segment HR-pQCT images of healthy distal radii [15,16]. The advantage of the active contouring algorithm is the inherent ability to pass over larger gaps in the bone. This is possible because the requirements for a contour to be both close to the object edges and smooth are balanced. Recently, three-dimensional morphological geodesic active contours (3D-GAC) have been implemented in Python, allowing for efficient determination of active contours on 3D images [17,18]. Hence, 3D-GAC appears to be a promising approach to create outer contours of HR-pQCT images of distal radius fractures.

The goal of present study was to investigate the use of fully automatic 3D-GAC to generate outer contours on HR-pQCT images of fractured radii. We validated the accuracy of these contours against hand-drawn contours using HR-pQCT images acquired throughout the first year of the healing process from 10 patients with fractured distal radii. We further assessed the robustness of the algorithm by comparing computed morphometric indices from intact distal radii HR-pQCT images of 73 subjects using contours generated from (i) the 3D-GAC and (ii) the scanner manufacturer’s default approach. Lastly, the reproducibility of the 3D-GAC based algorithm was assessed based on contours of 19 human subjects imaged six times using HR-pQCT.

## 2 Materials and Methods

### 2.1 Data

HR-pQCT images (XtremeCT II, 60.7 µm voxel size, 68 kV, 1470 µA, 43 ms integration time) were obtained from the database of a previous fracture study conducted at Innsbruck Medical University. Patients provided informed consent and participated in a study approved by the ethics committee of the Medical University of Innsbruck (UN 0374344/4.31). For each patient, scans of the fractured (up to 504 slices per scan distributed over three stacks) and contralateral (up to 168 slices per scan) radius were taken at six time points (1, 3, 5, 13, 26, and 52 weeks post-fracture) over the course of one year.

#### 2.1.1 Fractured Radii

Images of radius fractures for 10 out of the 75 patients that completed the study were selected from the database based on the highest available visual grading scores (VGS) [19] and well aligned stacks resulting in 60 HR-pQCT fractured radius images. For each image, the most distal slice before the appearance of the lunate fossa was identified, and all evaluations on these images were performed on slices proximal to that slice.

#### 2.1.2 Intact Radii

A total of 73 out of the 75 patients were selected, which had at least one HR-pQCT image of the contralateral, intact radius with a VGS of three or better, resulting in 438 images. Of these 73 patients, 19 had a VGS of three or better across all time points and were used for the reproducibility study.

### 2.2 Image Pre-Processing

Time-lapsed HR-pQCT images of each patient were registered using rigid image registration [20] to allow direct comparison of generated contours across all time points per patient.

### 2.3 Contours: Current Gold Standards

For ten randomly selected slices per fractured radius HR-pQCT image, hand-drawn 2D contours were generated by three researchers experienced in the processing of medical images (DCT, PRA, CJC), respectively, using the software of the scanner manufacturer. For this procedure, the 2D slice is visualised in the scanner manufacturer’s software, and using a computer mouse, the contour is drawn onto the image. The software provides zooming functionality and local edge detection to assist the operator in defining the outer contour. These hand-drawn 2D contours were used as the gold standard to validate the results of the 3D-GAC algorithm for fractured radii and to quantify the inherent inter-observer variability in hand-drawn contours of fractured radii.

For intact radii, the default manufacturer pipeline based on a 3D algorithm by Buie et al. [10], written in the Image Processing Language (IPL) of the scanner manufacturer (Scanco Medical AG, Switzerland), was used as the gold standard. All contours generated using the manufacturer’s software were exported as binary 3D images to h5 files [21], a hierarchical data format, for further processing using Python [22].

### 2.4 Novel Contouring Algorithm

The proposed algorithm uses 3D morphological geodesic active contours (3D-GAC) [17] implemented in the Python library Scikit-Image [23] using morphological operators [18]. The guiding principle for geodesic active contours is to minimize an energy term consisting of both internal and image energy. The internal energy penalizes non-smooth contours, while the image energy penalizes contours away from the voxels of interest. The image energy landscape, in this approach, is generated using an inverse Gaussian gradient, where the Gaussian blur removes local minima and the gradient is an operation to detect object edges.

The proposed contouring algorithm can be separated into four distinct steps. First, the image is segmented into sections that only contain one bone (here, radius or ulna). In a second step, an initial guess of the radius contour is generated. Third, the image is pre-processed and converted to an energy landscape. Finally, the 3D-GAC algorithm is applied in an iterative loop to isolate the surface of the radius.

#### 2.4.1 Separation of Radius and Ulna

3D-GAC are drawn to all intensity-based edges in an image and do not differentiate between objects of interest and other objects. Therefore, it is necessary to remove extraneous objects. Here, the ulna must be removed, as it is in close proximity to the radius. The watershed algorithm implemented in Scikit-Image [23] is used to separate these two bones (Figure 1), based on seed voxels of the different objects.

**Figure 1:**
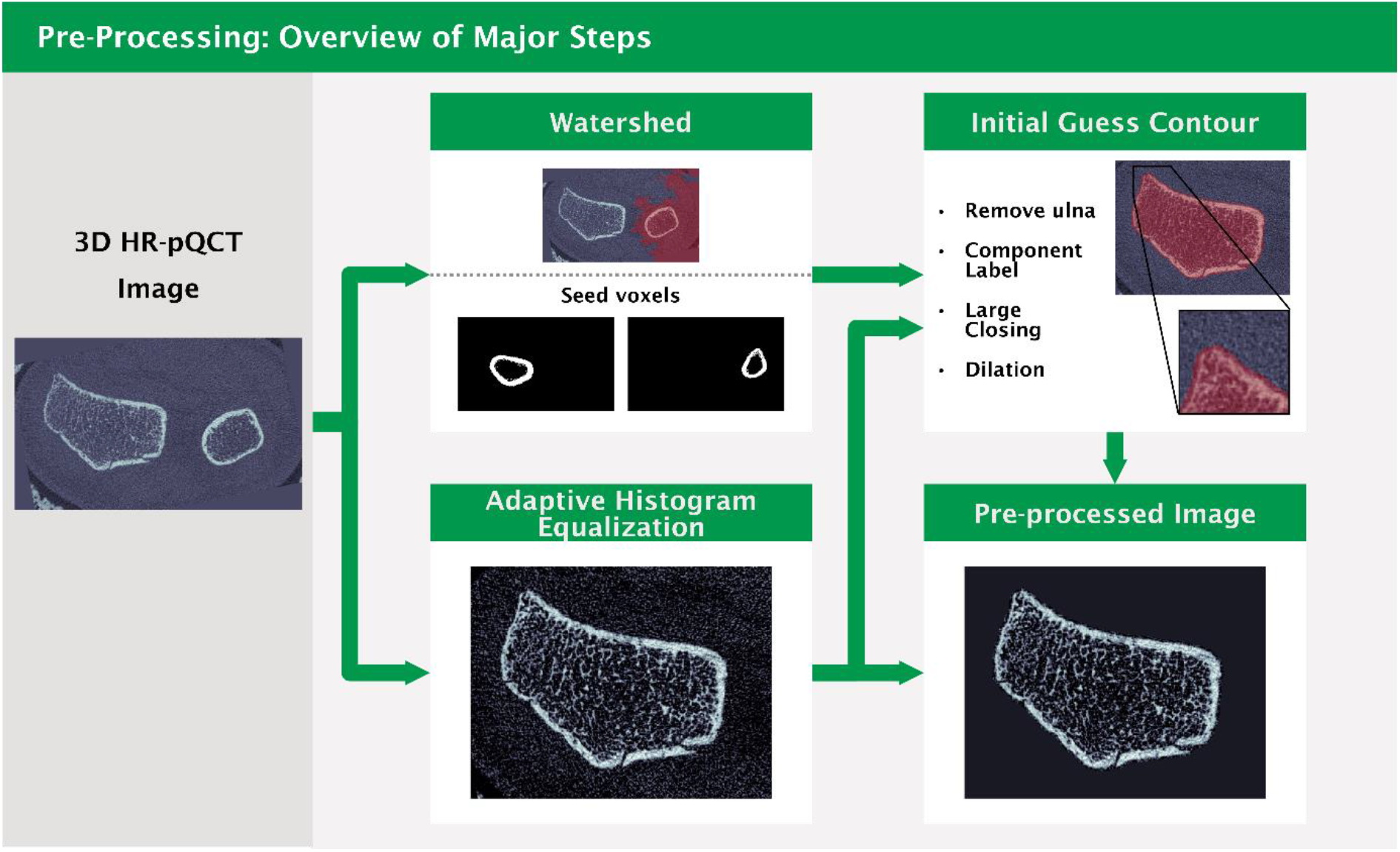
Overview of all pre-processing steps of the 3D-GAC approach. 3D high-resolution peripheral quantitative computed tomography (HR-pQCT) images (left) are coarsely segmented with the watershed algorithm using seed voxels of proximal slices (top middle). At the same time, adaptive histogram equalization is applied to the 3D HR-pQCT images to compensate for density differences in the proximal and distal cortex (bottom middle). The watershed segmentation and the equalized image are then combined using basic image processing tools to obtain an initial guess contour of the radius (top right). Finally, the initial guess mask is applied to the equalized image to obtain a clean image as input for the 3D-GAC algorithm (bottom right).

Since the radius and ulna are typically only a few voxels apart in ultra-distal scans, separating them using a global threshold has the risk of identifying both bones as a single object. Therefore, the radius and ulna are identified in a proximal slice (here, 35-39 slices from the most proximal slice to avoid end effects that may have been introduced as a result of image registration) using a threshold of 250 mg HA / cm^3^ and component labelling implemented in the Python library NumPy [24]. The largest component identified is the radius and the second largest one the ulna. Finally, a seeds image is generated by copying the labels of the radius and ulna in the proximal slice into the entire proximal half of a zeroed image (Figure 1). The distal part was left empty to avoid incorrect assignments of seeds in the distal region, where radius and ulna are not well separated.

The energy landscape for the watershed algorithm is generated from the HR-pQCT image by setting all negative values to zero and inverting the sign of all remaining elements. The algorithm is applied to the entire image such that every voxel – background or bone – is assigned to exactly either the radius or ulna (Figure 1).

#### 2.4.2 Initial Guess

To generate the initial guess contour for the 3D-GAC algorithm, the HR-pQCT image is first normalized to the range of zero to one. Afterwards, the image is equalized in the two orthogonal longitudinal planes independently using a contrast limited adaptive histogram equalization implemented in Scikit-Image [23], and the resulting two images are averaged. A threshold of 0.5 is applied and the ulna portion of the image identified during pre-processing is removed from the thresholded image. Finally, the largest connected component is extracted via component labelling, the structure is closed with a closing distance of 50 voxels, the remaining interior holes are filled in an additional component-labelling step and the structure is dilated five iterations to obtain the initial guess (Figure 1).

#### 2.4.3 Pre-Processing of Image

To prepare the image from the scanner for input to the 3D-GAC algorithm, the image is first cropped to the bounds of the initial guess contour to reduce the image size, which reduces computational cost. The image is then normalized to the range of zero to one and equalized in the same way as for the creation of the initial guess. The catchment basins from the pre-processing watershed algorithm are used to set those voxels not belonging to the radius section to zero. Finally, the voxels outside the initial guess are set to the mean value of the initial guess surface voxels, which are those voxels that make up the perimeter of the initial guess. The final pre-processed image (Figure 1) is then contoured using the 3D-GAC algorithm.

#### 2.4.4 3D Geodesic Active Contours (3D-GAC)

The 3D-GAC algorithm takes an initial guess, an energy landscape and outputs an optimized contour. Since the algorithm is easily trapped in local minima, i.e. a contour that does not match the surface of the radius, an iterative application of 3D-GAC to energy landscapes containing increasing levels of detail is part of our approach. For each iteration, the pre-processed image is used to create an energy landscape (Figure 2). For this, the image is first padded in the longitudinal direction by 20 voxels at each end using the NumPy [24] edge mirroring setting. Gaussian blurring is then applied to smooth the energy landscape (Figure 2). Finally, the padding voxels are removed. Since finer details of the radius can get lost when Gaussian blurring with a large sigma, three iterations of 3D-GAC are run using decreasing sigmas (14.0, 3.0, 1.5 voxels) to allow for an iterative approximation of the true radius contour (Figure 2). Note that for the first iteration, image resolutions are halved after blurring to speed up computations. Each 3D-GAC application is then run using the default parameters of SciPy [25] with the number of iterations set to five (Figure 2). For the first iteration of 3D-GAC the initial guess (Figure 1) is used while successive iterations then use the output of the previous iteration of the 3D GAC algorithm as the new initial guess (Figure 2). As a final step, component labelling [24] followed by a selection of everything not connected to the background is performed to remove holes in the contour that can appear at the distal image boundary.. The optimized final contour can then be used to segment the radius from the HR-pQCT image background for further analysis.

**Figure 2:**
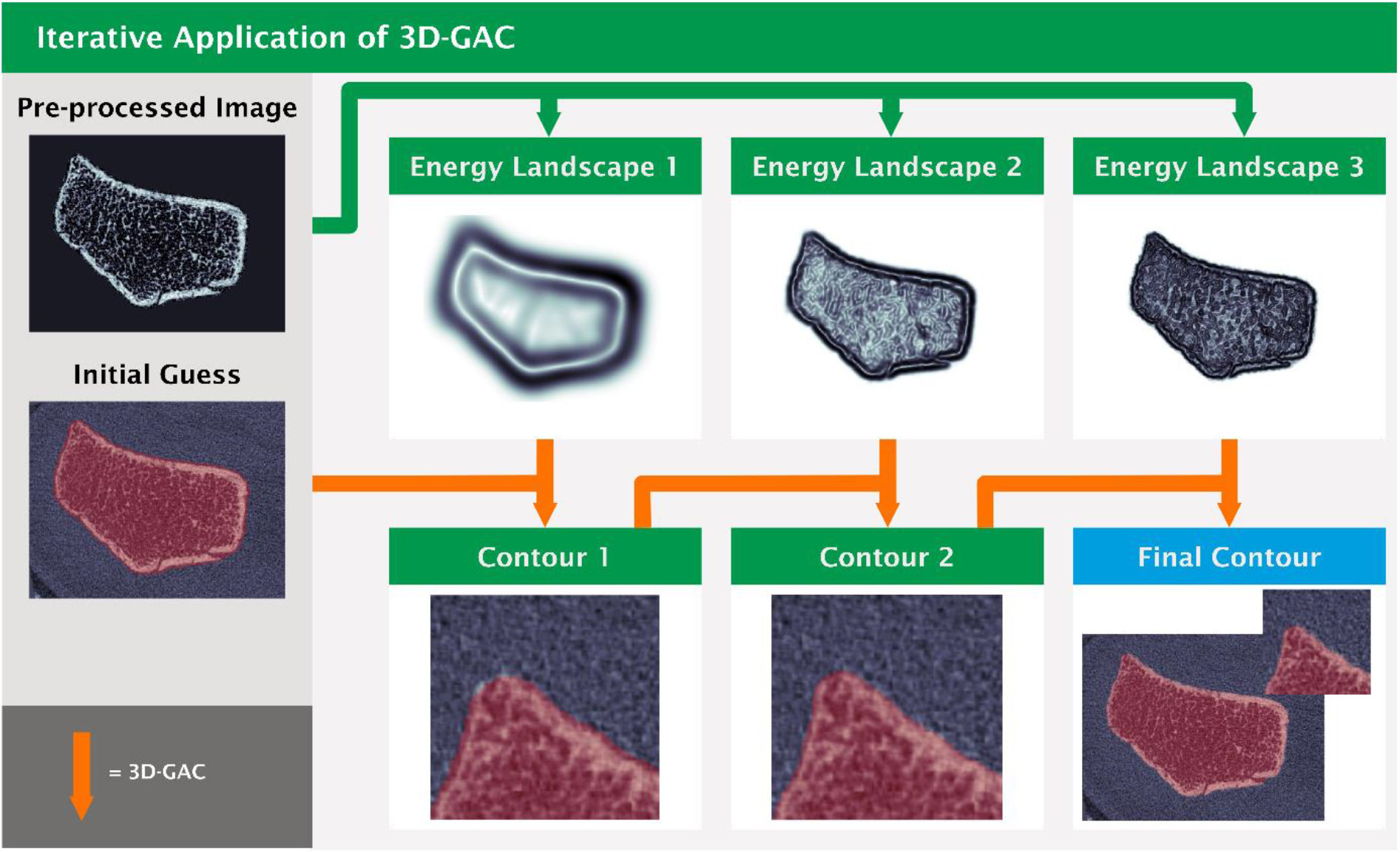
Iterative approximation of the final contour via multiple 3D-GAC applications. The pre-processed images (top left and Figure 1) is converted to an energy landscape using Gaussian blurring and computing the gradient to detect edges. Per iteration the blurring is reduced (top left to right) to allow for an increased amount of detail to be picked up by the 3D GAC per iteration. For the first iteration of 3D GAC the initial guess (bottom left and from Figure 1) is used. Successive iterations then use the output of the previous iteration of the 3D GAC algorithm as the new initial guess (bottom).

### 2.5 Morphometric Indices

The Scanco system provides the standard patient evaluation script for XtremeCT II devices, which is the gold standard approach of generating cortical and trabecular masks [12] and computing bone morphometric indices. Total volumetric bone mineral density (Tt.vBMD), trabecular volumetric bone mineral density (Tb.vBMD), cortical volumetric bone mineral density (Ct.vBMD), trabecular area (Tb.Ar), cortical area (Ct.Ar), trabecular bone volume fraction (BV/TV), trabecular number (Tb.N), trabecular thickness (Tb.Th), trabecular separation (Tb.Sp), cortical perimeter (Ct.Pm), and cortical thickness (Ct.Th) were computed for this study. The default setup uses outer contours generated by the approach of Buie et al. as input [10]. However, the standard patient evaluation pipeline also accepts alternative outer contours as input. Outer contours generated from our novel 3D-GAC method were used as input to the standard patient evaluations and the resultant morphometrics were compared to those of the manufacturer’s default pipeline.

### 2.6 Study Design

To assess the quality of the generated contours, we compared them to the respective gold standard for two different datasets (fractured and intact distal radii).

#### 2.6.1 Fractured Radii: Comparison of the 3D-GAC Approach to Hand-Drawn Contours

The 3D-GAC method was run for all fracture images. Contours were visually categorised as acceptable, mistakenly including parts of the ulna, having obvious missing parts, or having both of these issues. Only images categorised as acceptable were used for additional quantitative analysis steps. Agreement between hand-drawn contour operators was assessed by computing the Dice score between their respective 2D contours for each image. The Dice score is able to quantify differences in automatically generated contours against reference contours yielding values in the range from zero to one with zero indicating no overlap while one indicates a perfect match. The Dice score is close to percent agreement when comparing contours that are similar to each other. Additionally, the distance transform of the area between the edges of 2D contours between operators was computed. Accuracy of the 3D-GAC approach was determined by taking the median Dice score between the automatic and the three hand-drawn 2D contours for each of the slices per image. Furthermore, all hand-drawn 2D contours created by the three operators were combined to yield a smallest and largest contour. The voxels between these two contours describes an area of uncertainty for the hand-drawn contours. Voxels in the smallest hand-drawn 2D contour but not in the automatic one and voxels in the 3D-GAC contour but not in the largest hand-drawn 2D contour were extracted and a distance transform was applied as a measure for deviations of the 3D-GAC contours from the hand-drawn 2D ones. The automated default manufacturer pipeline was also run on all fracture images to generate outer contours and categorisation as for the 3D-GAC contours was performed.

#### 2.6.2 Intact Radii: Comparison of the 3D-GAC and Manufacturer’s Approach

The 3D-GAC approach and the default manufacturer pipeline were run on all intact radii. Contours of both approaches were visually assessed and categorized as described previously for the fractured radii. For each patient, one image for which both approaches were successful was randomly chosen for further analysis. Accuracy of the 3D-GAC method was assessed by computing the Dice score between each contour of the 3D-GAC method and the respective contour of the manufacturer’s method. Additionally, the distance transform of the volume between the surfaces of both approaches was computed as another measure of how much the two methods deviated from each other. Reproducibility was determined by computing the Dice score of each follow-up image with the baseline image for each patient. Morphometrics were computed to assess how strongly the computation of standard morphometric indices is influenced by the choice of outer contour.

### 2.7 Statistics

Bland-Altman plots were created in Python to assess systematic differences between hand-drawn and 3D-GAC contours. Additionally, Bland-Altman plots were generated to assess systematic deviations between morphometric indices generated from the default manufacturer and 3D-GAC contours. To allow visual comparison of the magnitudes of deviations between contour area and different morphometric parameters, all Bland-Altman plots had their y-axis scaled by 13% of the maximum x-value throughout this paper, which was the maximum value not leading to clipping of data. Robust linear models were fitted for Bland-Altman plots with Huber’s T for M estimation using the statsmodels package in Python [26]. This package also provides two-sided Student’s t-tests for the linear models with zero slope and intercept of each Bland-Altman plot being the null-hypotheses, respectively. Here, the default significance level of p < 0.05 was implemented.

Statistical significance for the comparison of morphometrics was performed using a paired Student’s t-test. Significance level was set to p<0.05.

## 3 Results

### 3.1 Fractured Radii: Comparison of the 3D-GAC Approach to Hand-Drawn Contours

From the 60 images, 20 were successfully contoured by both 3D-GAC and the default manufacturer pipeline. The default manufacturer pipeline failed 40 times, while the 3D-GAC approach failed only for four images (Figure 3a). Typically, the default manufacturer pipeline failed due to missing parts of the radius as a result of fracture gaps in the cortex (N=33), inclusion of parts of the ulna in the contour (N=2) or both (N=5). The 3D-GAC approach failed due to missing parts of the radius (N=2), inclusion of parts of the ulna in the contour (N=1), or inclusion of parts of the background in the contour (N=1). Furthermore, the default manufacturer pipeline only showed a high success rate for images taken one-year post fracture (Figure 3b), whereas the 3D-GAC worked similarly well for all time points. Note that the default manufacturer pipeline has not been designed to handle images of fractured radii and therefore one could expect the unfavourable outcome. Visually, contours matched the given bone structures well within the inter-operator variability (Figure 4). Agreement was found between the 3D-GAC approach and hand-drawn contours with a mean Dice score of 0.992 ± 0.004. This is within the range of inter-operator agreement: (operator 1 vs operator 2) 0.993 ± 0.005, (operator 1 vs operator 3) 0.993 ± 0.006, and (operator 2 vs operator 3) 0.994 ± 0.005. Evaluating the six appointments individually, Dice scores were high (>0.985), though some increase in the inter-quartile range of the Dice scores was observed between three and thirteen weeks post fracture (Figure 3c). This coincided with a decrease in contouring success (Figure 3b). For voxels of the 3D-GAC deviating from the area of uncertainty, 82% of these contour voxels were only one voxel away from at least one hand-drawn contour. This was within the inter-observer variability, as we observed (operator 1 vs operator 2) 57%, (operator 1 vs operator 3) 60%, and (operator 2 vs operator 3) 58% of deviations being less than one voxel away between operators (Figure 3d). Comparing the contour areas between the 3D-GAC method and the three operators, agreement was high with errors being: -1.14 ± 0.76%, -0.40 ± 0.62%, and -0.22 ± 0.73% for each operator, respectively. An underestimation of the total area was observed for the 3D-GAC contours (Figure 5). This underestimation (Figure 5d) is less than the measured difference comparing operator 1 to operator 3 (Figure 5b): slope (−0.003 vs -0.011) and intercept (−0.076 vs 0.090 (both not significant)).

**Figure 3:**
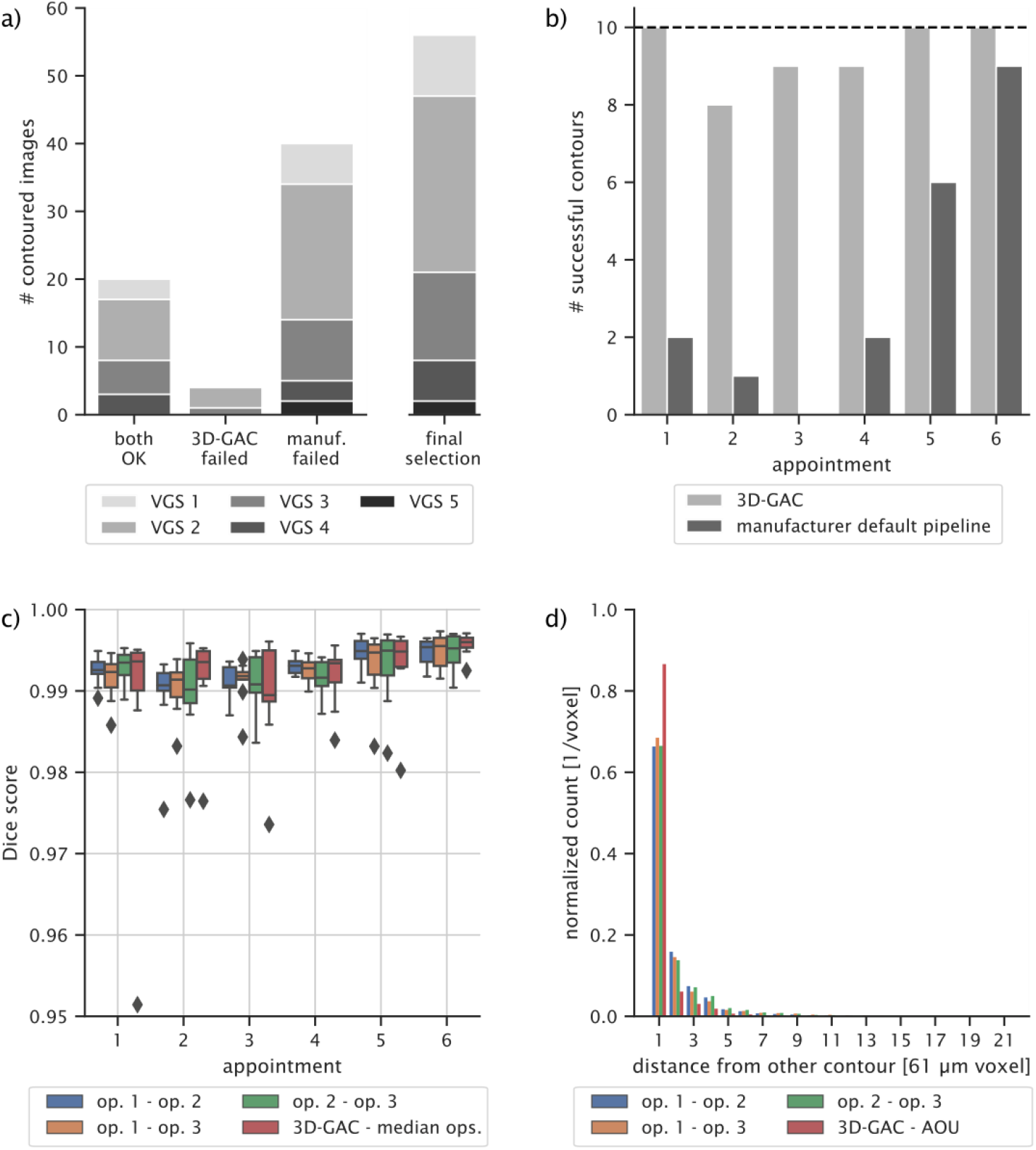
Accuracy of the 3D-GAC approach vs hand-drawn contours for scans of fractured radii. a) Histogram of successful and failed contours for the 3D-GAC and the default manufacturer pipeline. The final bar shows the images selected for further analysis. Note that this pipeline was not designed to handle scans of fractured radii. b) Histogram of successful contours for the two approaches per patient time-appointment. Dashed line indicates 100% success. c) Dice scores computed for all images between the hand-drawn slices of all operators (op) respectively. The corresponding slices of the 3D-GAC contours were also compared against the hand-drawn slices, respectively, and the average median Dice score is shown. d) Histogram of the distance of all voxels different between the 3D-GAC and the area of uncertainty (AOU) for the hand-drawn contours of each operator, respectively. The AOU is defined as the area between the contour obtained by only accepting voxels present in all hand-drawn contours and the contour obtained by accepting voxels present in at least one hand-drawn contour.

**Figure 4:**
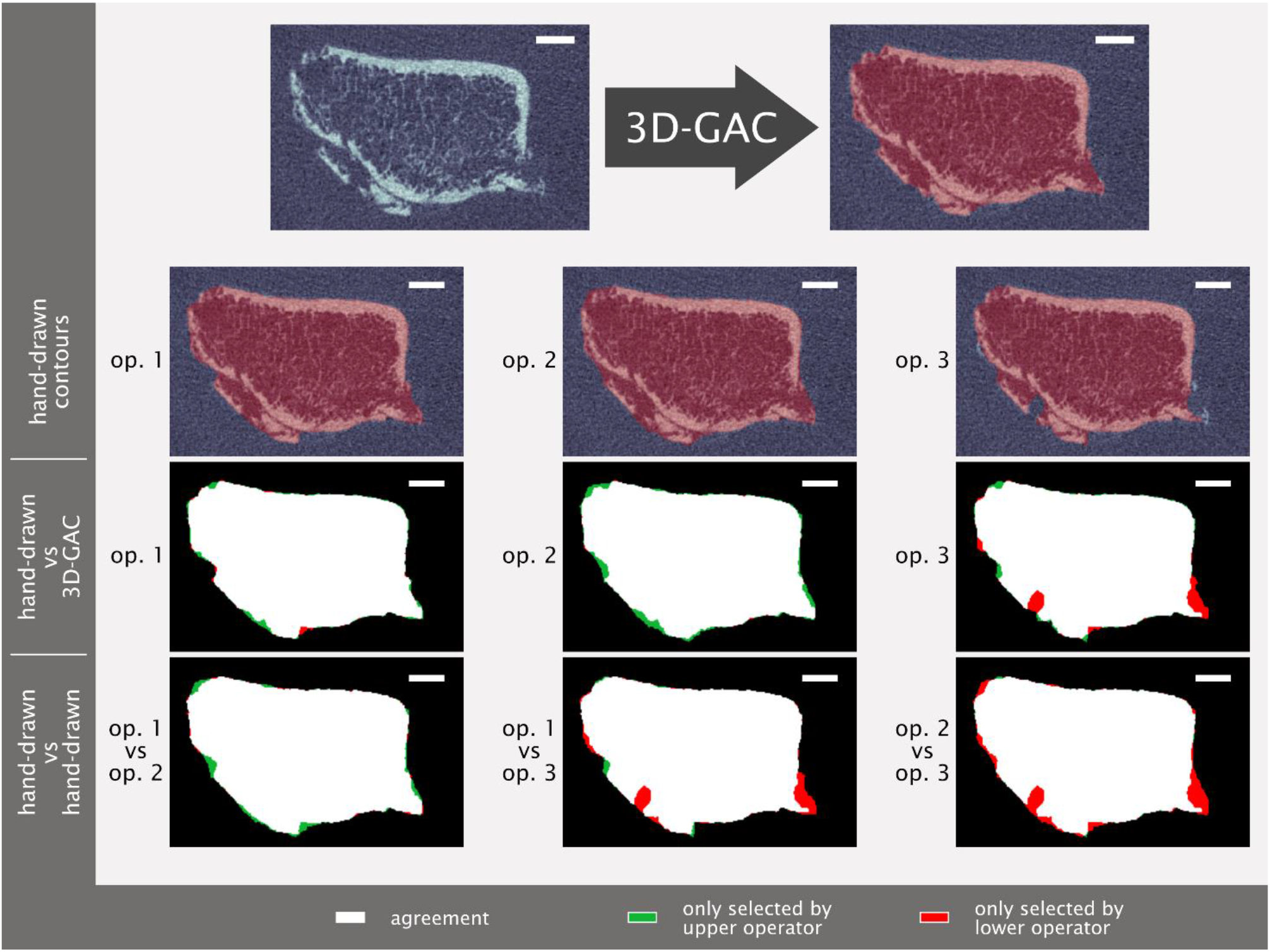
Visual comparison of hand-drawn contours (by three different operators (op)) and 3D geodesic active contour (3D-GAC) generated contours for a representative challenging slice from one week post-fracture. Visual differences between the 3D-GAC and hand-drawn contours are similar to those between hand-drawn contours of different operators. Scale bar is 2 mm.

**Figure 5:**
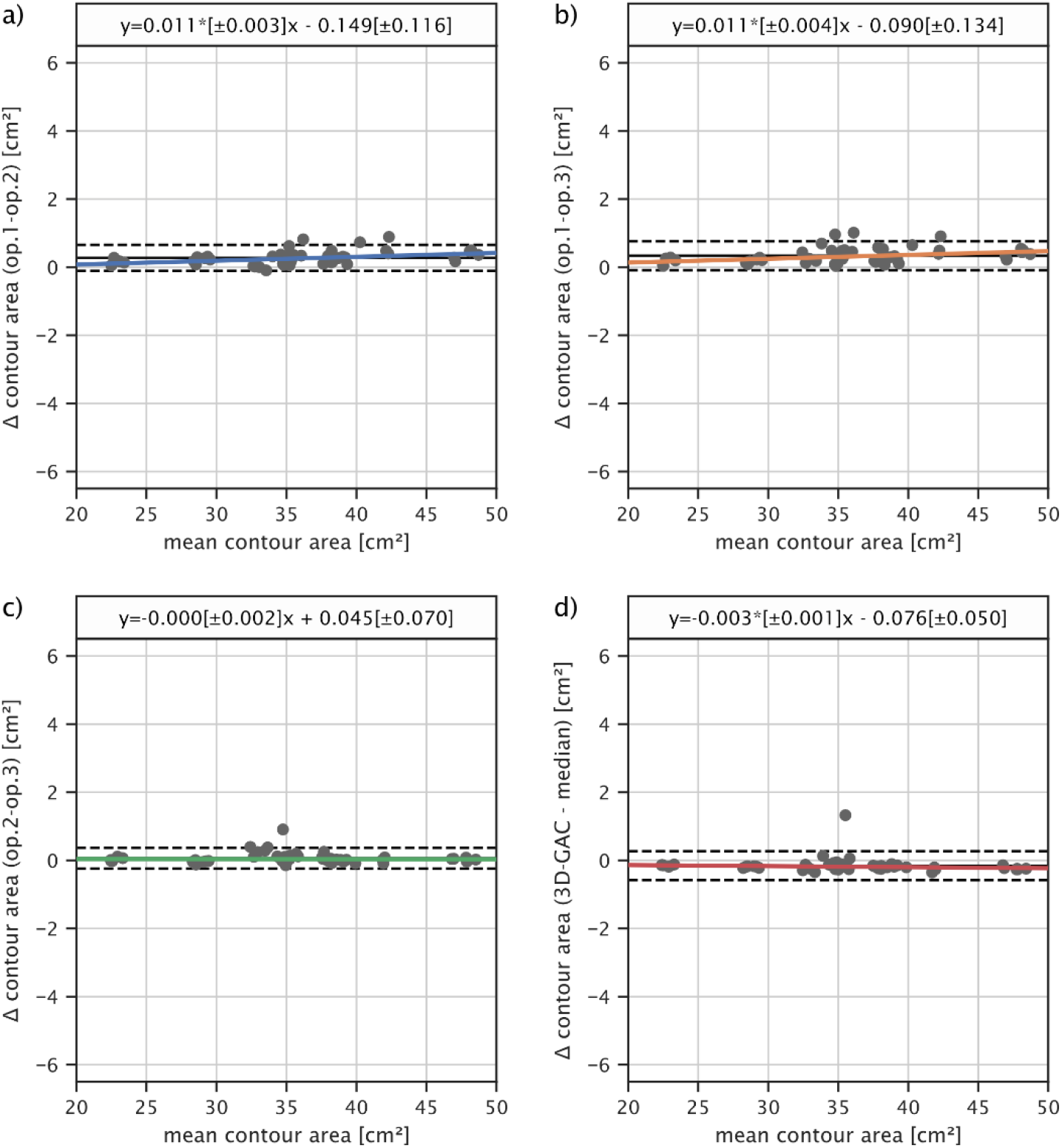
Bland-Altman plots of the mean contour area per image. a) Comparing operator 1 (op. 1) to operator 2 (op. 2). b) Comparing operator 1 to operator 3 (op. 3). c) Comparing operator 2 to operator 3. d) Comparing the automatic 3D-GAC contours with the median (computed per slice) of all three operators. Area computed with the 3D-GAC approach agrees within the observable inter-operator variability with the area computed from hand-draw contours.

### 3.2 Intact Radii: Comparison of the 3D-GAC and Manufacturer’s Approach

From the 438 images of intact radii, 341 were successfully contoured by both approaches. The default manufacturer pipeline failed for 85 images, the 3D-GAC approach failed 3 times, and both approaches failed 9 times (Figure 6a). The default manufacturer pipeline failed due to the inclusion of parts of the ulna into the radius contour (N=42), omission of large parts of the radius (N=37) (Figure 7), inclusion of parts of the background in the contour (N=5), or inclusion of parts of the ulna into the radius contour while omitting large parts of the radius (N=10). The 3D-GAC approach failed due to the inclusion of background (N=8), omission of large parts of the radius (N=1), omission of parts of the radius and inclusion of parts of the ulna in the contour (N=2), and all mentioned failure modes combined (N=1). The VGS for the failed contours of the manufacturer’s approach ranged from one to five (N=7 for VGS 1, N=14 for VGS 2, N=16 for VGS 3, N=30 for VGS 4, N=27 for VGS 5). For the thirteen contours that only failed for the 3D-GAC approach (N=1 for VGS 2, N=5 for VGS 4, N=6 for VGS 5).

**Figure 6:**
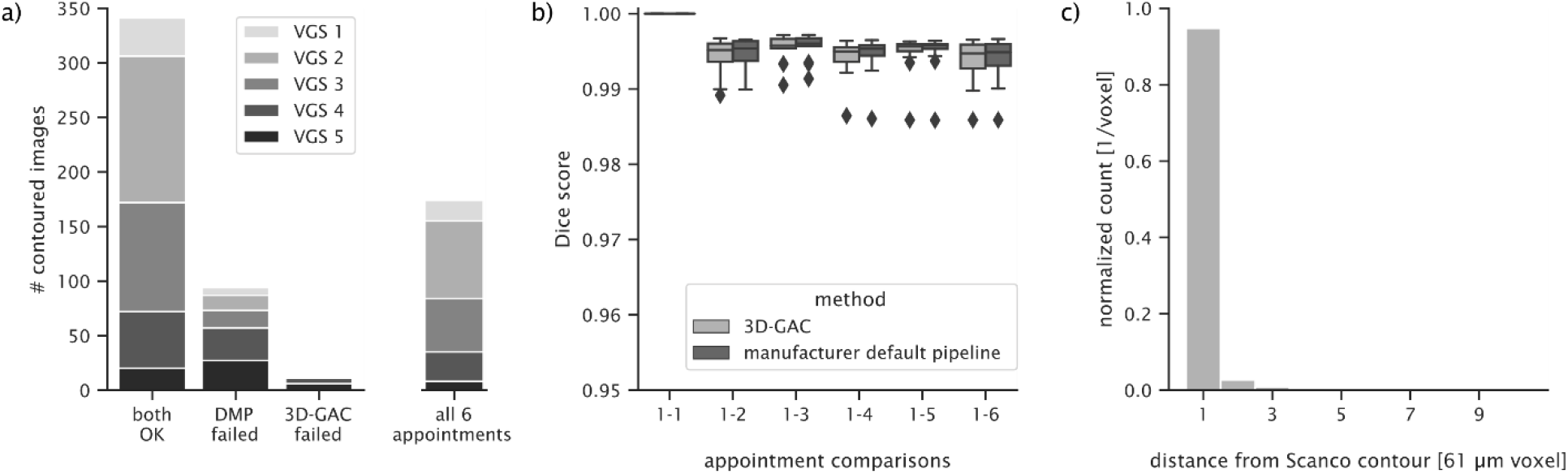
Accuracy and reproducibility of the 3D-GAC vs the default manufacturer pipeline (DMP). a) Histogram of successful and failed contours for the 3D-GAC and the DMP with indicated image quality. The final bar shows the images selected for the reproducibility analysis. b) Dice scores of contours of all time points relative to the initial scan for both approaches show the reproducibility of each method. c) Histogram of the distance of all voxels different between the 3D-GAC and the DMP contour from the respective DMP contour.

**Figure 7:**
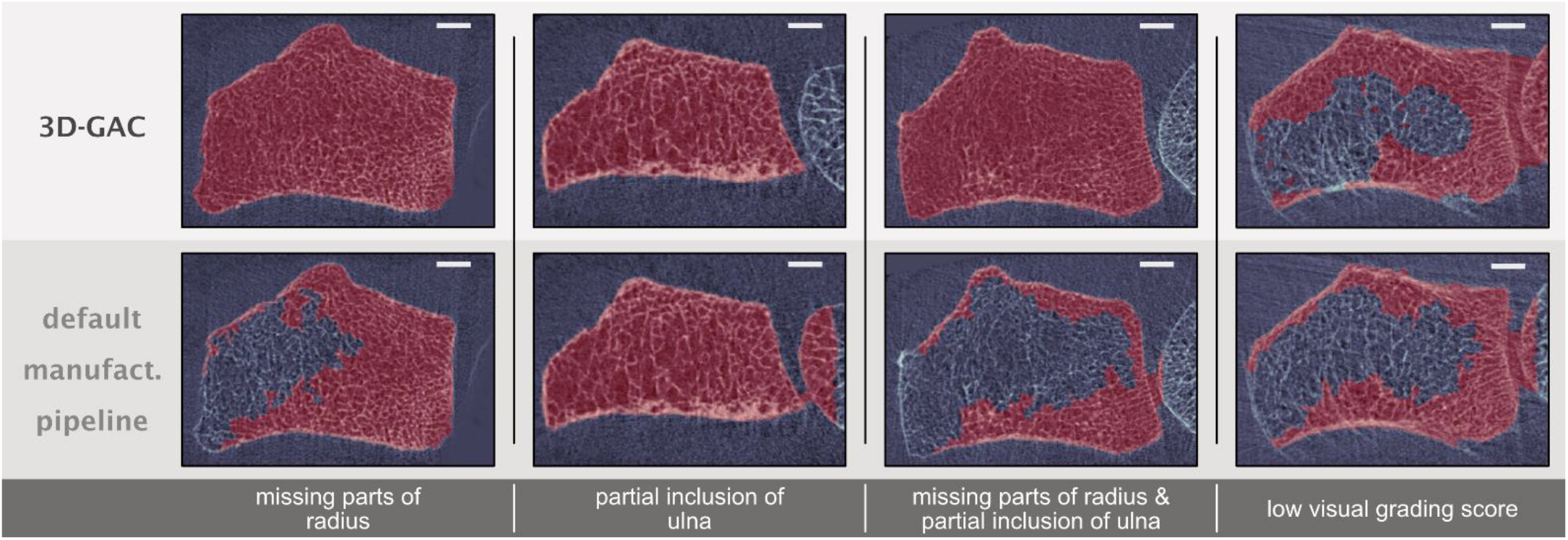
Examples of failure modes when contouring distal radii. Contours from the 3D-GAC and the default manufacturer pipeline are compared for four challenging contouring cases: A thin cortex leading to missing parts of the radius, the ulna (partly being visible on the right) being close to the radius (bone on the left) and hence being included in the contour, the combination of the previous two, and a low visual grading score. Only the 3D-GAC approach manages to generate visually correct contours in most cases. Scale bar is 2 mm.

Of the 341 images successfully contoured by both approaches, one image per patient (N = 72, one patient from the initial dataset was not successfully contoured by both approaches) with sufficient image quality (VGS of three or better) was randomly selected for comparative analysis. Overall agreement between the contours generated by both approaches was high with a mean Dice score of 0.996 ± 0.001, with typical Dice score values for contours with missing parts of the radius being around 0.85. Reproducibility, measured for the 19 patients indicated in the materials section, was on the same level as the default manufacturer pipeline, wherein agreement with the baseline image for both methods and for all appointments was found (median Dice score > 0.994 for all time-points) (Figure 6b). For those voxels that were either part of the 3D-GAC contour and not part of the contour from the default manufacturer pipeline or vice versa, 91% were at most one voxel away from the manufacturer’s contour (Figure 6c). Bland-Atman plots revealed agreement in morphometric indices with a slight thickness- and density-based bias in cortical parameters and no bias in trabecular parameters (Figure 6). Mean deviations were below 1% for all parameters except for Ct.Th (1.341%) (Table 1).

**Table 1:** Deviations of the bone morphometrics derived using 3D-GAC based contours compared to the manufacturer’s default pipeline.

| Morphometric Index | Deviation from Gold Standard |
| --- | --- |
| Tt.vBMD | $0.190 \pm 3.668\%$ |
| Tb.vBMD | $0.662 \pm 3.226\%$ |
| Ct.vBMD | $-0.131 \pm 2.437\%$ |
| Tb.Ar | $0.510 \pm 4.920\%$ |
| Ct.Ar | $-0.710^* \pm 2.530\%$ |
| BV/TV | $0.540 \pm 2.830\%$ |
| Tb.N | $-0.093 \pm 1.497\%$ |
| Tb.Th | $0.147 \pm 1.017\%$ |
| Tb.Sp | $0.181 \pm 1.625\%$ |
| Ct.Pm | $-0.333 \pm 2.402\%$ |
| Ct.Th | $-1.341^* \pm 4.911\%$ |
| **Morphometric Indices: total volumetric bone mineral density (Tt.vBMD), trabecular volumetric bone mineral density (Tb.vBMD), cortical volumetric bone mineral density (Ct.vBMD), trabecular area (Tb.Ar), cortical area (Ct.Ar), trabecular bone volume fraction (BV/TV), trabecular number (Tb.N), trabecular thickness (Tb.Th), trabecular separation (Tb.Sp), cortical perimeter (Ct.Pm), cortical thickness (Ct.Th).**<br>**\*indicates significant deviation ($P < 0.05$)** |  |

## 4 Discussion

### 4.1 Fractured Radii: Comparison of the 3D-GAC Approach to Hand-Drawn Contours

One of the main aims of the present study was to develop a fully automatic outer contouring approach for HR-PQCT images of distal radius fractures. An image processing pipeline using a combination of adaptive histogram equalization, watershed segmentation, iterative Gaussian blurred energy landscapes, and 3D-GAC was established such that 56 out of 60 images were automatically contoured successfully. In our hand-contoured reference dataset, we observed clear inter-operator variability, which is a known issue of hand-contouring bone [27]. On average, the automatically generated contours by the 3D-GAC algorithm agreed well with operators. Interestingly, both hand-drawn and 3d-GAC contours had slightly larger inter-quartile ranges for appointments 2 and 3, likely as a result of the formation of a callus in this time period and the resulting low mineralization of background voxels visible in the HR-pQCT images. This makes it harder for both operators and the algorithm to decide on the correct contour. Given that no systematic deviations were detected between operators and the 3D-GAC approach and that deviations from individual operators were within the range of inter-operator variability (Figure 5), the 3D-GAC contouring approach developed in this study provides accurate contours for HR-pQCT images of fractured distal radii compared to the gold standard.

As a point of reference, the default manufacturer’s pipeline was used to generate outer contours of the fractured images. The default manufacturer pipeline only reliably contoured HR-pQCT images taken one-year post fracture (Figure 3b). As this pipeline was not designed to work with images of impaired cortices, this might indicate structural weaknesses in the cortex of the healing radii even 6 months post fracture. This observation is in agreement with recent findings suggesting that fracture repair is a long-term process which might take as long as two years [28].

### 4.2 Intact Radii: Comparison of the 3D-GAC and Manufacturer’s Approach

The second aim of the present study was to evaluate the robustness of the 3D-GAC contours against the default manufacturer pipeline by comparing the resultant morphometrics. Notably, the failure rate of the automated manufacturer default pipeline was much higher than that of the 3D-GAC approach for the HR-pQCT images of the intact radii (94 vs 12 failed contours, respectively). This may be a result of the patient cohort used in the present study since fracture patients have been shown to have inherently poorer bone quality than standard populations [29]. One of the major reasons for failure of the default manufacturer pipeline was the inclusion of the ulna into the outer contour of the radius, which is a known issue of this approach that is implementation dependent [10]. The second reason for failure was having obvious missing parts, which appeared to happen when the cortex had very thin or low-density structures (Figure 7). The authors would like to emphasise that these failures of the default manufacturer pipeline can be, and often are, overcome by generating hand-drawn contours using a built-in tool of the manufacturer’s software. However, as noted previously, the cost in person-hours of such a solution is prohibitive in large patient cohort studies. An automated solution for this issue was proposed by Buie and co-workers [10]; here, they increased the number of dilation steps of the manufacturer’s pipeline but noted that this often lead to unwanted smoothing of the contour [10]. Moreover, defining per-image numbers of dilation steps would reduce the degree of automation drastically. Both approaches were affected by image quality, as expected. However, the 3D-GAC approach only failed for one image with sufficient image quality to be considered for inclusion in HR-pQCT studies [8,27,30]. Trabecular morphometric indices generated using the successful 3d-GAC contours agreed well with those generated using the default manufacturer pipeline with deviations well below the precision error of these indices [31].

Interestingly, for the cortical parameters, accuracy was slightly worse for Ct.Th generated using the 3D-GAC approach, showing deviations of 1-2%; however, these differences were still less than 1 voxel in thickness and thus at the accuracy limit of the device (Figure 6). Furthermore, differences in cortical thickness were at the reproducibility limit of the standard patient evaluation (1.3-3.9% dependent on subject age) [32] and were still below differences observed between patient groups from the literature, e.g. healthy and osteopenic women (12.8%) [33]. Given the improved success rate and accurate determination of morphometric indices, the 3D-GAC approach showed an overall improvement in robustness compared to the current gold standard.

**Figure 6:**
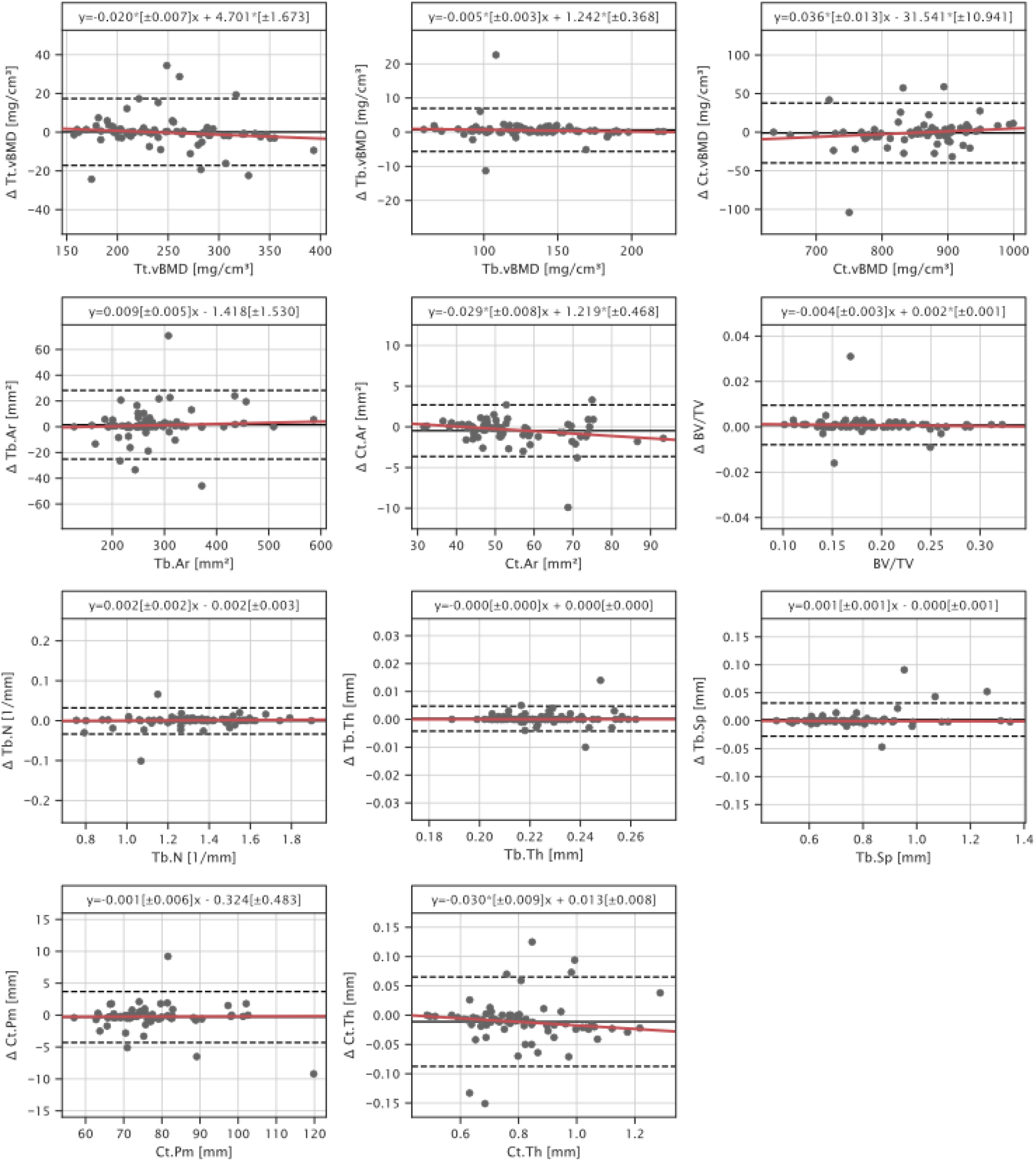
Bland-Altman plots of morphometric indices characterize potential differences in 3D-GAC and contours from the default manufacturer pipeline. Robust linear model fits are shown in red. Regression parameters are presented above each plot with slope and intercept parameters statistically significantly different from zero being indicated by (*, P<0.05). Indices are total volumetric bone mineral density (Tt.vBMD), trabecular volumetric bone mineral density (Tb.vBMD), cortical volumetric bone mineral density (Ct.vBMD), trabecular area (Tb.Ar), cortical area (Ct.Ar), trabecular bone volume fraction (BV/TV), trabecular number (Tb.N), trabecular thickness (Tb.Th), trabecular separation (Tb.Sp), cortical perimeter (Ct.Pm), cortical thickness (Ct.Th).

Lastly, the present study aimed to determine the reproducibility of the 3D-GAC approach using sequential image scans of the intact radii taken over the course of a year. Here, agreement with the baseline contour was found to be high for all time points (Dice scores > 0.994). Again, deviations in Ct.Th were highest, but well within the reproducibility limit of the standard patient evaluation.

Given the improved robustness and sufficient reproducibility, the 3D-GAC approach appears to be a viable option for clinical studies of both fractured and non-fractured radii.

### 4.3 Limitations

This study is not without limitations. The repeat scans of the contralateral site were spaced over a year, and thus may include variations in bone due to natural remodelling. Additional reproducibility scans could not be taken for these patients due to radiation concerns. Future studies should include follow-up scans at protracted intervals to assess how sensitive the 3D-GAC approach is to local and global remodelling. Due to the large structural changes happening during fracture repair, the images of the fracture site could not be regarded as repeat scans. Therefore, no direct assessment of reproducibility was possible for images of fractured radii. However, the high accuracy observed between the hand-drawn and 3D-GAC contours of the fractured images indicates a high level of reproducibility. Another limitation is the low number of hand-drawn contour slices (270) generated to use as reference. While time was a limiting factor (operators reporting to take roughly 1-2 minutes per slice) for creating more hand-drawn contours, we compensated by including images of the intact contralateral site to cover a large variety of bone morphologies in our assessment of the 3D-GAC robustness and repeatability. These images of the intact radius provided a means to benchmark the 3D-GAC approach against existing automatic approaches for intact radii. Note that the 3D-GAC contours were generated only proximal to the lunate fossa, excluding the articulating surface. Although this limits the range over which the 3D-GAC approach can be applied, the morphometric analysis of the standard patient evaluation has only been developed for that region; as such, this region remains the most relevant for clinical studies. Lastly, no hand-corrected contours were generated for intact radii to estimate the bias introduced by the automated manufacturer and 3D-GAC protocols. However, recent studies have already done this for the gold standard [34] and, given the good agreement of the 3D-GAC approach with the gold standard, the requirement for minor manual corrections should be on a similar level.

## 5 Conclusion

Our proposed 3D-GAC algorithm provides a unified pipeline for generating outer contours from HR-pQCT images of distal radii, including both fractured and intact bones. Importantly, our method automatically contours radii both accurately and reproducibly, while showing robustness when dealing with a wide variety of cortex structures. This will facilitate future clinical studies using large patient cohorts to support the development of improved treatment protocols of radius fractures.

## 6 Acknowledgements

The authors acknowledge the Swiss National Science Foundation (320030L_170205), the European Union’s Horizon 2020 research and innovation programme under the Marie Skłodowska -Curie grant (agreement 841316), and the ETH Postdoctoral Fellowship for financial support. The funders had no role in study design, data collection and analysis, decision to publish, or preparation of the manuscript.

